# Allosteric inhibition of the SARS-CoV-2 main protease – insights from mass spectrometry-based assays

**DOI:** 10.1101/2020.07.29.226761

**Authors:** Tarick J. El-Baba, Corinne A. Lutomski, Anastassia L. Kantsadi, Tika R. Malla, Tobias John, Victor Mikhailov, Jani R. Bolla, Christopher J. Schofield, Nicole Zitzmann, Ioannis Vakonakis, Carol V. Robinson

## Abstract

Following translation of the SARS-CoV-2 RNA genome into two viral polypeptides, the main protease M^pro^ cleaves at eleven sites to release non-structural proteins required for viral replication. M^Pro^ is an attractive target for antiviral therapies to combat the coronavirus-2019 disease (COVID-19). Here, we have used native mass spectrometry (MS) to characterize the functional unit of M^pro^. Analysis of the monomer-dimer equilibria reveals a dissociation constant of *K*_d_ = 0.14 ± 0.03 μM, revealing M^Pro^ has a strong preference to dimerize in solution. Developing an MS-based kinetic assay we then characterized substrate turnover rates by following temporal changes in the enzyme-substrate complexes, which are effectively “flash-frozen” as they transition from solution to the gas phase. We screened small molecules, that bind distant from the active site, for their ability to modulate activity. These compounds, including one proposed to disrupt the catalytically active dimer, slow the rate of substrate processing by ~35%. This information was readily obtained and, together with analysis of the x-ray crystal structures of these enzyme-small molecule complexes, provides a starting point for the development of more potent molecules that allosterically regulate M^Pro^ activity.

---

The coronavirus, SARS-CoV-2, is the etiological agent of the 2020 pandemic that has claimed >550,000 lives and affected more than 12 million people as of July 2020.^[1]^ Coronaviruses have long existed in nature and have made zoonotic transmission to humans. Despite the tragic and widespread effects of these sudden occurrences, we do not yet have validated anti-viral treatments targeting coronavirus infections. SARS-CoV-2 packages a large RNA genome of ~30k bases, two-thirds of which encodes for two polyproteins (pp1a and pp1b). These polyproteins are processed into 15 non-structural proteins (nsps) that are liberated from the long polypeptide chains by two viral proteases, the papain-like protease (nsp 3) and the 3C like protease (nsp 5). The latter species, named the main protease M^pro^ is a cysteine protease that cleaves the viral polyproteins at eleven sites to generate twelve non-structural proteins (nsp4-nsp15). Included in these nsps are those involved in the replication machinery (e.g., the RNA-dependent RNA polymerase nsp12).^[2]^ Inhibition of M^pro^ impairs the ability of the virus to replicate.

Analogous to the 2004 SARS-CoV main protease, the functional unit of the SARS-CoV-2 M^Pro^ is a homodimer (Figure 1).^[3]^ Several encouraging strategies to inhibit M^Pro^ have been explored via its covalent inhibition.^[3a, 3b]^ Non-covalent modulators, including compounds that disrupt the dimer interface, have not been investigated to the same extent. Drugs that bind non-covalently can often be fine-tuned to diffuse through membranes and bind to target proteins with high affinity.^[4]^ Moreover, these species lack reactive warheads often leading to higher chemical stability than their covalent counterparts and a reduction in undesirable toxic effects due to irreversible binding to host proteins and nucleic acids. Therefore, non-covalent compounds serve as a promising means to inhibit viral proliferation.

**Figure 1.**
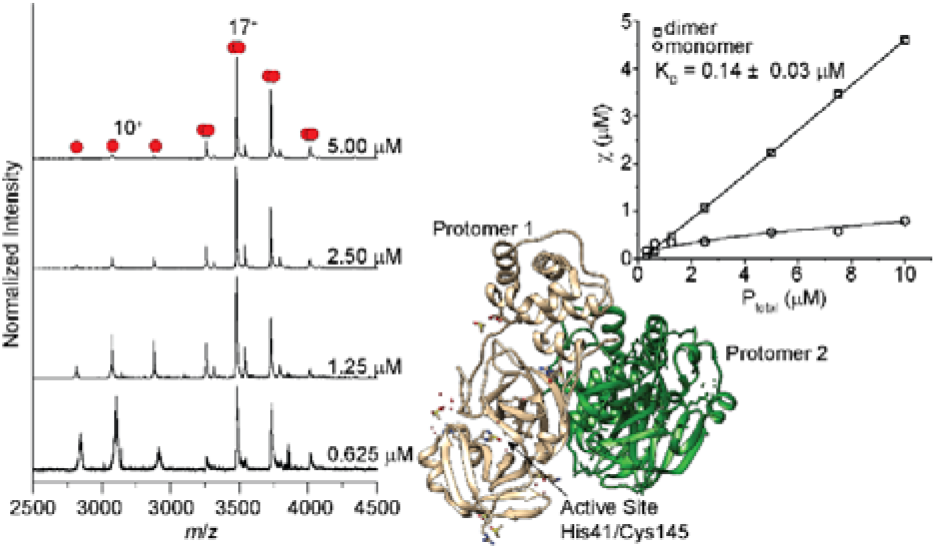
Analysis of M^Pro^ by native MS. (left) Native mass spectra for M^Pro^ at different concentrations. (right) Representative plot of mole fraction versus concentration to quantify the dissociation constant and a view from an x-ray structure of unligated M^Pro^ (PDB 6YB7).

Previous studies of the SARS-CoV M^Pro^ dimer identified nano to micromolar dissociation constants.^[5]^ We report the dissociation constant for SARS-CoV-2 M^Pro^ determined using native mass spectrometry (MS), which can directly identify and quantify the relative amounts of the oligomeric state of a protein in solution.^[6]^ To probe the monomer-dimer equilibrium we recorded native mass spectra over a range of M^Pro^ concentrations from 0.313 to 10.0 μM in an aqueous buffer containing 200 mM ammonium acetate (pH = 7.4). At a protein concentration of 5 μM, two well-resolved charge state distributions are readily identified. The high-abundant signals centered at the 17^+^ charge state correspond to a deconvoluted mass of 67,591 ± 0.5 Da, consistent with the expected sequence mass of dimeric M^Pro^ (67,592 Da). A minor peak series between m/z ~2750 and 3500 Th, is also observed. These signals, centered at a 10^+^ charge state, correspond to a deconvoluted mass of 33,795 ± 3 Da, in excellent agreement with the mass of monomeric M^Pro^ with wildtype N- and C-termini (33,796 Da). As the concentration of protein is decreased in a series of stepwise dilutions from 5.0 μM to 0.625 μM, the signals corresponding to the M^Pro^ monomer increase concurrent with a decrease in the peak intensities assigned to the dimer.

To extract a monomer-dimer equilibrium constant we performed measurements in triplicate at seven different protein concentrations (from 0.325 to 10 μM) and plotted the mole fraction of each species as a function of total protein concentration (Figure 1). Excellent agreement was found between the measured values and a monomer-dimer equilibrium binding model (see Supporting Information) providing high confidence in the derived dissociation constant, *K*_d_ = 0.14 ± 0.03 μM. We conclude that, analogous to SARS-CoV M^Pro^, the SARS-CoV-2 variant has a high propensity to form dimers.

Following the deposition of high-resolution SARS-CoV-2 M^Pro^ structures to the protein databank,^[3a, 3b]^ a fragment screen was released providing structural insight towards 96 candidates that bind M^Pro [3c]^ Three of these candidates were identified as binding to the dimer interface; 23 non-covalent and 48 covalent hits were found to bind the active site. We focused our study to those small molecules that bind non-covalently to M^Pro^ and considered their potential to both destabilize the dimer and modulate substrate cleavage. We identified four candidates – three that bind to the solvent exposed surface (x0390, x0425, and x0464) and one that binds within the dimer interface (x1187). We incubated each species with 5 μM M^Pro^ at a ligand concentration range of 1 to 100 μM (0.125 – 20-fold molar excess). Following incubation for 30 mins we recorded mass spectra to investigate their effect on the monomer-dimer equilibrium. x0390, x0425, and x0464 showed no appreciable perturbation of the monomer-dimer ratios (see Supporting Information). However, the effect of x1187 on dimerization was clearly apparent (Figure 2). In the absence of x1187, under the solvent conditions used to dissolve the small molecules, the charge state distribution is centered at 15^+^ reinforcing the notion that M^Pro^ is predominantly a dimer under a range of solution conditions. Following the addition of a 10-fold molar excess of x1187 the peaks assigned to monomers, centered at 10^+^ charge state, are observed to increase. After addition of a 20-fold molar excess of x1187 the fractional abundance of the monomer peaks increases further to ~15%.

**Figure 2.**
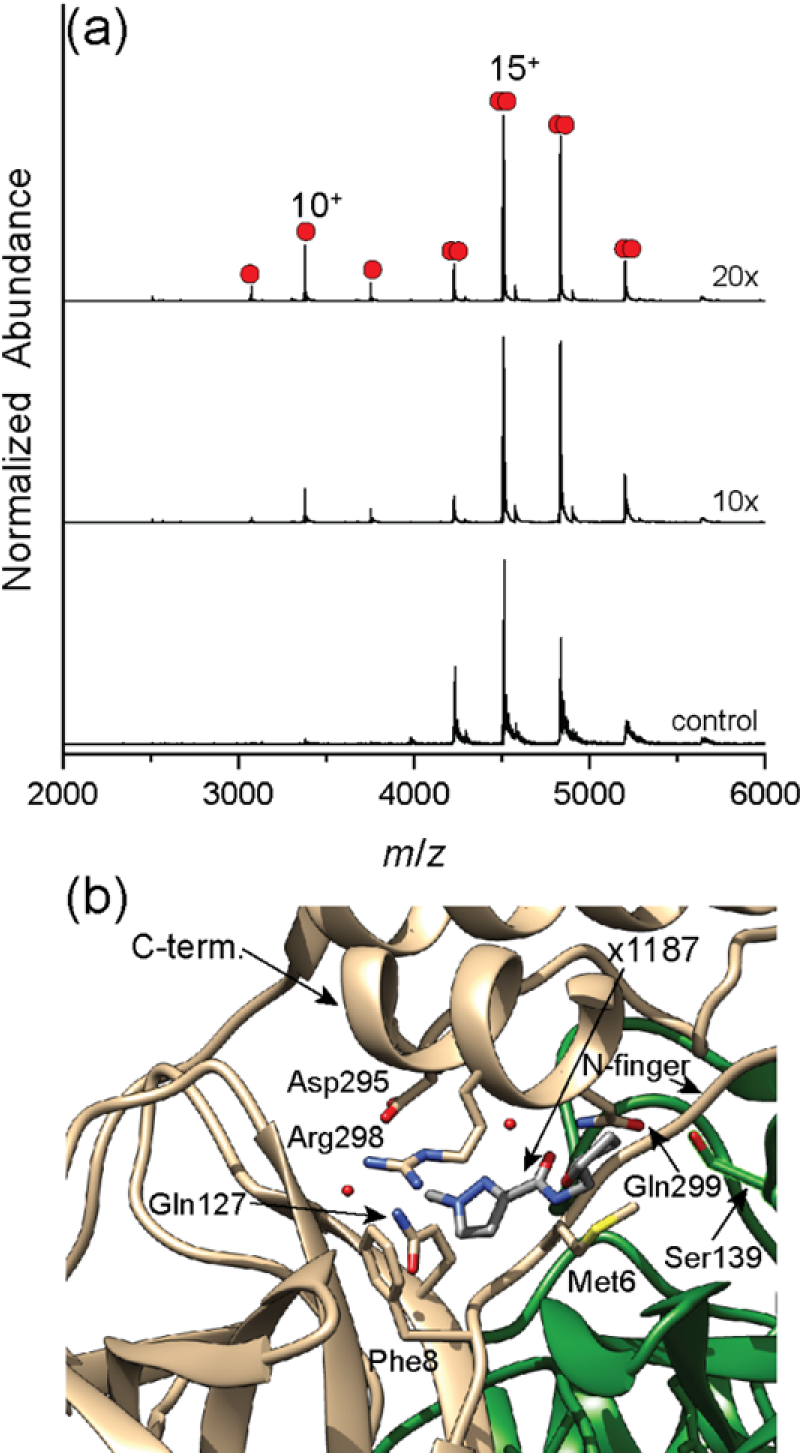
(a) Native mass spectra of 5 μM M^Pro^ with the addition of different molar equivalents of x1187. To maintain similar solution conditions to the samples containing x1187, the control contains 10% DMSO. (b) Detailed view of dimer interface where x1187 binds (PDB 5FRA).

Inspection of the crystal structure identifies the likely origin of this destabilization; x1187 rests across the dimer interface, packing into a hydrophobic pocket partially comprised of key residues Met6 and Phe8 on the N-finger (Figure 2). The long axis of x1187 rests along the C-terminal helix of protomer 1 and is proximal to Ser139 on protomer 2, a critical residue for proper function and assembly.^[7]^ The position of x1187 is important as it rests in a pocket crucial to formation of a dimer with a productive topology,^[8]^ prompting us to consider whether or not this, or the other small molecules that bind non-covalently, mediate catalysis.

To identify the ability for these molecules to mediate the proteolytic activity of M^Pro^ we first characterized the substrate turnover in the absence of small molecules using an MS-based kinetic assay. This assay quantifies the change in observable enzyme-substrate complexes as a function of time. These complexes are notoriously difficult to isolate and quantify as they are turned over rapidly; but as they transition from solution to the gas phase, they are effectively “flash-frozen”. Quenching the reaction enables interrogation of these transient species captured in the MS instrument.^[9]^ A mass spectrum collected 30 s after initiating the cleavage reaction is shown (Figure 3). Several satellite peaks are observed alongside the main charge state series, one of which corresponds to the enzyme substrate complex with a mass of 68,781 ± 2 Da, in agreement with dimeric M^Pro^ bound to a single 11-mer substrate (1192 Da). We also identified a series of peaks with a deconvoluted mass of 68,188 ± 0.5 Da, consistent with the acyl-enzyme complex that is formed by reaction of the nucleophilic Cys145 with the scissile Q-S peptide bond of the substrate to give a covalent TSAVLQ-enzyme complex of mass 68,192 Da (Figure 3). A summary of the M^Pro^-substrate complexes we identified are listed in Table 1.

**Table 1.** M^Pro^ and M^Pro^-substrate complexes identified by native MS

| Species | Measured Mass (Da)<br>[a] | Expected Mass (Da) | Δmass (Da) |
| --- | --- | --- | --- |
| $M^{\text{Pro}}$ (Monomer) | $33,795 \pm 3$ | 33,796 | -1 |
| $M^{\text{Pro}}$ (Dimer) | $67,591 \pm 0.5$ | 67,592 | -1 |
| Enzyme-substrate complex | $68,781 \pm 2$ | 68,784 | -2 |
| Enzyme-TSAVLQ complex | $68,188 \pm 0.5$ | 68,192 | -4 |

[a] Uncertainty in deconvoluted mass determined using at least three charge states.

**Figure 3.**
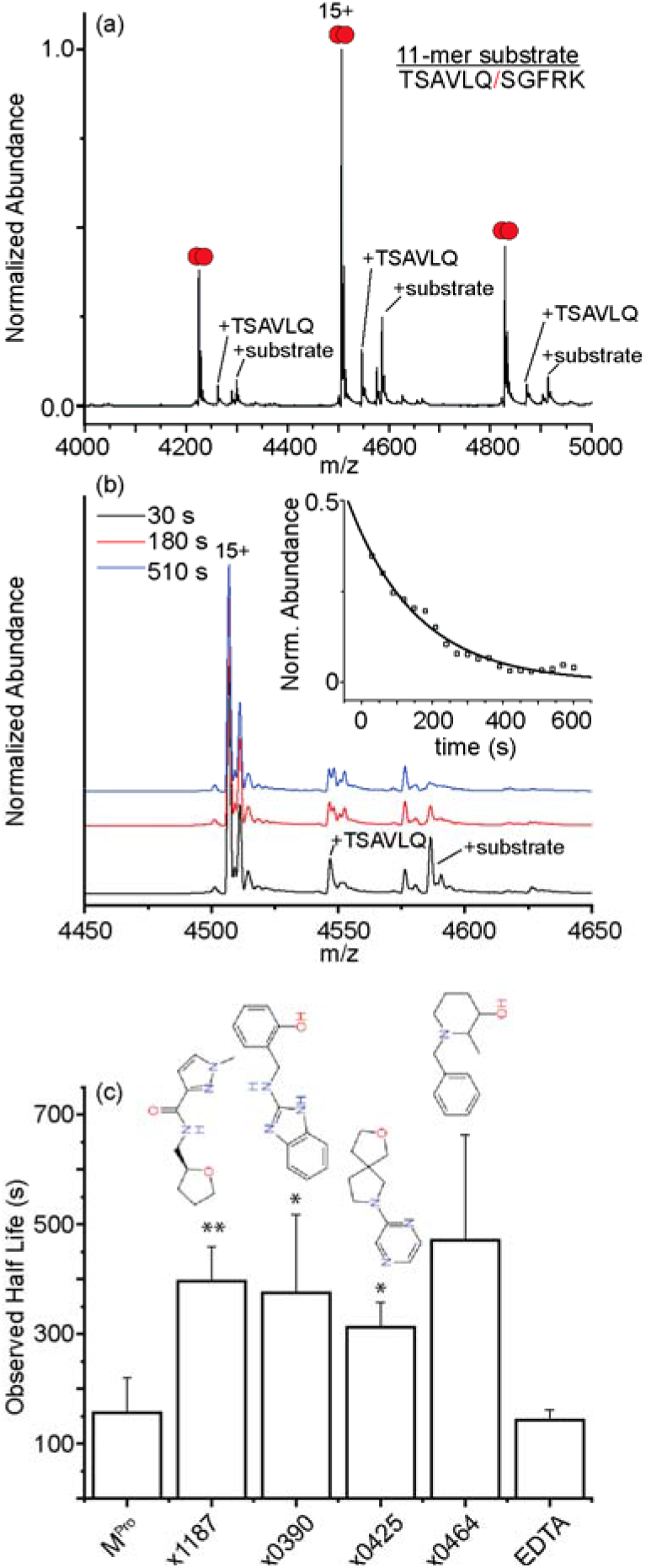
(a) Native mass spectrum for 5 μM M^Pro^ with 50 μM of the 11-mer substrate at t = 30 s. Peaks labelled TSAVLQ and +substrate indicate acyl-enzyme complex and the non-covalent enzyme-substrate complex respectively. (b) mass spectra for the 15^+^ charge state at three representative times along the substrate cleavage reaction. Inset shows a plot of the relative abundance of the enzyme-substrate complex as a function of time. Solid line indicates the fit to a unimolecular kinetics model. (c) Bar plot summarizing half-lives of the enzyme-substrate complex in the presence of different small molecules. Error bars represent standard deviation (n=3 independent replicates).*p<0.05, **p<0.001 (to M^Pro^ values). Representative mass spectra and kinetic plots for each dataset are shown in the Supporting Information.

Interestingly, we did not observe any peaks corresponding to monomeric M^Pro^ bound to the 11-mer substrate or acyl-enzyme complex (see Supporting Information). We considered several reasons for the absence of signals for substrate-bound monomers. One possibility is that the monomers have enhanced activity compared to the dimer. The timescale we measure for substrate cleavage by the dimer is on the order of minutes. Thus, if monomers were to bind and process substrates the monomer activity would need to be enhanced by ~100-fold (complete turnover by ~6 s), otherwise we would readily capture monomer-substrate complexes. This is unlikely, as other assays have indicated that monomeric M^Pro^ is inactive.^[5c]^ We surmise that monomers are not only inactive but also they do not bind the 11-mer substrate with high affinity.

An expansion of the 15^+^ charge state at three representative time points reveals that as time evolves, the signal for the enzyme-substrate complex at 4586 *m*/*z* is depleted (Figure 3). At our longest timepoint (10 min), the signals for the enzyme-substrate complex are no longer present in detectable quantities indicating that by this time the substrate 11-mer has been proteolyzed and the products released (see Supporting Information). A plot of the relative abundance of the decay of the enzyme-substrate complex versus reaction time reveals that the decay is exponential (Figure 3). The experimental data can be readily fit to a single-step unimolecular kinetics model, providing a relatively straightforward means to compare half-life changes in the presence of inhibitors. A plot comparing the half-lives obtained by incorporating the 20-fold molar excess of the different small molecules into the reaction mixtures is shown (Figure 3 and Supporting Information). We note that the presence of these small molecules increases the lifetime of the enzyme-substrate complex, in some cases by as much as ~38%.

As control experiments, we first characterized the propensity for substrate turnover using a potent inhibitor IPA3 that covalently modified Cys145 and found no appreciable changes in substrate turnover, formation of the enzyme-substrate complex, or acyl-enzyme complex (See Supporting Information). We also characterized the kinetic behavior of M^Pro^ in the presence of 50-fold molar excess of EDTA. We found no appreciable change in the depletion of the enzyme-substrate complex, suggesting that trace amounts of divalent metal cations, (e.g., Zn^2+^) when added in isolation, are not modulators of M^Pro^ activity.^[10]^

All of the small molecule candidates tested inhibit M^Pro^ proteolytic activity and bind to locations distant from the active site

(Figure 4). Specifically, x1187 binds at the dimer interface and x0390, x0425, and x0464 bind to solvent-exposed pockets remote form the active site. While x1187 disrupts the dimer, its proximity to key residues in the vicinity of the N-finger may also allow it to intervene with the N-finger in regulating activity^[8]^ and allows us to conclude that all of these fragments inhibit substrate cleavage via allosteric regulation.

**Figure 4.**
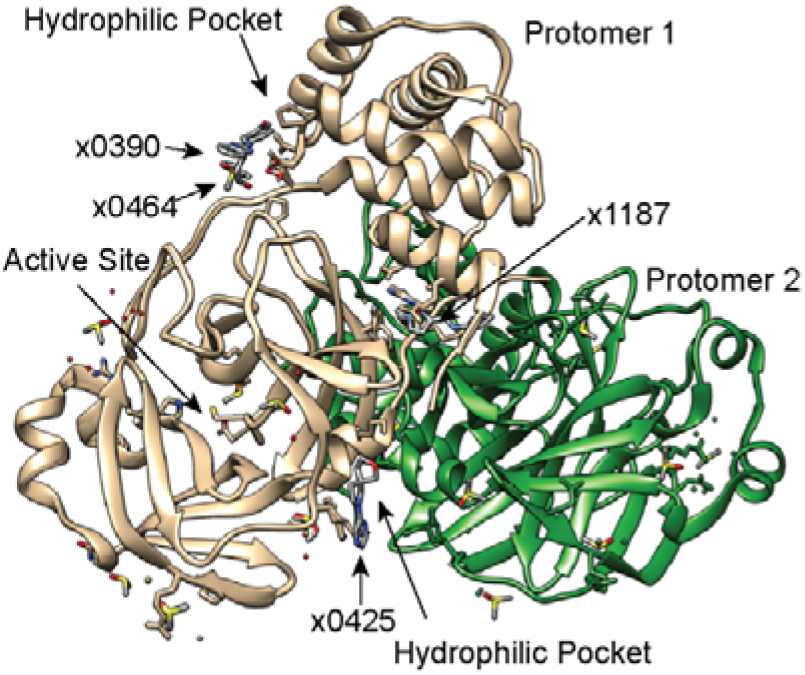
Superimposition of views of x-ray structures for unligated M^Pro^ (PDB 6YB7) and M^Pro^ bound to x1187 (PDB 5FRA), x0390 (PDB 5REC), x0464 (PDB 5REE), and x0425 (5RGJ).

The absence of monomeric M^Pro^ bound to substrate raises an interesting question – how is pp1a/pp1ab processed upon viral infection? With a high propensity to dimerize, we speculate that viral polypeptide processing by M^Pro^ is carried out through a pp1a/pp1ab oligomer rather than M^Pro^ monomers. Approximating the volume of a mammalian cell to be 2.43 × 10^−3^ nL,^[11]^ our findings indicate that only ~200,000 molecules of M^Pro^ are required for the enzyme to preferentially form catalytically active dimers and process viral polypeptides. Moreover, the monomer-dimer equilibrium may be influenced by other cellular factors; the assembly state may shift to favor dimers given that M^Pro^ could be enriched subcellularly through the formation of endosomal vesicles.^[12]^ This may support the formation of pp1a/pp1ab dimers that transiently associate, facilitating the release of M^Pro^ from within the viral polypeptides.^[5c]^ Regardless of the mechanisms behind the initial steps of M^Pro^ maturation, our findings provide evidence that disruption of the catalytically active dimer with small molecules provides an opportunity to explore potent antivirals with low risk for toxicity.

More generally, the therapeutic potential of these, or analogous small molecules, can be readily assessed through the dual MS approaches described here: probing the monomer-dimer equilibrium and measuring efficacy in reducing substrate turnover rates. These relatively straightforward measurements delineate the mechanism of action of these allosteric regulators - either via disruption of the dimer interface or reversible non-competitive binding, distal from the active site. Overall, in addition to highlighting new ways of assaying these small molecules, the results provide a starting point for lead optimization chemistry with the ultimate goal of deactivating SARS-CoV-2 M^Pro^ and reducing the burden of the ongoing COVID-19 pandemic.

## Acknowledgements

We are grateful to Claire Strain-Damerell, Petra Lukacik, David Owen and Martin A. Walsh for providing the M^Pro^ expression construct and advise on protein production, and to the Diamond Light Source XChem facility, led by Frank von Delft, for providing small molecule ligands. Work in the C.V.R. laboratory is supported by a Medical Research Council (MRC) program grant (MR/N020413/1), a European Research Council Advanced Grant ENABLE (695511), and a Wellcome Trust Investigator Award (104633/Z/14/Z). T.J.E is supported by the Royal Society as a Royal Society Newton International Fellow. C.A.L. is supported by the European Commission as a Marie Skłodowska Curie Postdoctoral Fellow. A.L.K was supported by the Oxford Glycobiology Institute Endowment. TRM thanks Biotechnology and Biological Sciences Research Council (BB/M011224/1). TJ was supported by the Oxford-GSK-Crick Doctoral Program in Chemical Biology, EPSRC (EP/R512060/1) and GlaxoSmithKline. C.J.S. thanks the Wellcome Trust and the Medical Research Council for support. We are grateful for generous support provided by the University of Oxford COVID-19 Research Response fund and its donors (BRD00230).

## Entry for the Table of Contents

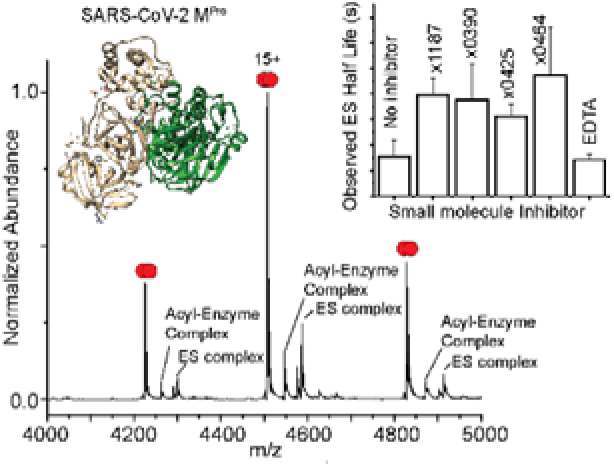
The SARS-CoV-2 main protease monomer-dimer equilibrium has been characterized using native mass spectrometry. A straightforward MS-based kinetic assay that quantifies the changes in the amounts of enzyme-substrate complex with time was used to capture M^Pro^ protease activity. The presence of several small molecules that bind non-covalently to M^Pro^ and do not compete for the active site slow the processing of the substrate, providing a straightforward means for prioritizing and optimizing potential antivirals.

## Notes

### Competing Interest Statement

The authors have declared no competing interest.

## References

[1] Values taken from: https://coronavirus.jhu.edu

[2] Y. Gao, L. Yan, Y. Huang, F. Liu, Y. Zhao, L. Cao, T. Wang, Q. Sun, Z. Ming, L. Zhang, J. Ge, L. Zheng, Y. Zhang, H. Wang, Y. Zhu, C. Zhu, T. Hu, T. Hua, B. Zhang, X. Yang, J. Li, H. Yang, Z. Liu, W. Xu, L. W. Guddat, Q. Wang, Z. Lou, Z. Rao, Science 2020, 368, 779–782.

[3] Z. Jin, X. Du, Y. Xu, Y. Deng, M. Liu, Y. Zhao, B. Zhang, X. Li, L. Zhang, C. Peng, Y. Duan, J. Yu, L. Wang, K. Yang, F. Liu, R. Jiang, X. Yang, T. You, X. Liu, X. Yang, F. Bai, H. Liu, X. Liu, L. W. Guddat, W. Xu, G. Xiao, C. Qin, Z. Shi, H. Jiang, Z. Rao, H. Yang, Nature 2020, 582, 289–293;

4. L. Zhang, D. Lin, X. Sun, U. Curth, C. Drosten, L. Sauerhering, S. Becker, K. Rox, R. Hilgenfeld, Science 2020;

5. A. Douangamath, D. Fearon, P. Gehrtz, T. Krojer, P. Lukacik, C. D. Owen, E. Resnick, C. Strain-Damerell, A. Aimon, P. Ábrányi-Balogh, J. Brandaõ-Neto, A. Carbery, G. Davison, A. Dias, T. D. Downes, L. Dunnett, M. Fairhead, J. D. Firth, S. P. Jones, A. Keely, G. M. Keserü, H. F. Klein, M. P. Martin, M. E. M. Noble, P. O’Brien, A. Powell, R. Reddi, R. Skyner, M. Snee, M. J. Waring, C. Wild, N. London, F. von Delft, M. A. Walsh, 2020.

[4] S. Zhang, X. Xue, L. Zhang, L. Zhang, Z. Liu, Chemical Biology & Drug Design 2015, 86, 1411–1424.

[5] W.-C. Hsu, H.-C. Chang, C.-Y. Chou, P.-J. Tsai, P.-I. Lin, G.-G. Chang, Journal of Biological Chemistry 2005, 280, 22741–22748;

8. V. Graziano, W. J. McGrath, L. Yang, W. F. Mangel, Biochemistry 2006, 45, 14632–14641;

9. B. Xia, X. Kang, Protein & Cell 2011, 2, 282–290.

[6] K. Gupta, J. A. C. Donlan, J. T. S. Hopper, P. Uzdavinys, M. Landreh, W. B. Struwe, D. Drew, A. J. Baldwin, P. J. Stansfeld, C. V. Robinson, Nature 2017, 541, 421–424.

[7] S. Chen, J. Zhang, T. Hu, K. Chen, H. Jiang, X. Shen, Journal of Biochemistry 2007, 143, 525–536.

[8] N. Zhong, S. Zhang, P. Zou, J. Chen, X. Kang, Z. Li, C. Liang, C. Jin, B. Xia, Journal of Virology 2008, 82, 4227–4234;

13. X. Xue, H. Yang, W. Shen, Q. Zhao, J. Li, K. Yang, C. Chen, Y. Jin, M. Bartlam, Z. Rao, Journal of Molecular Biology 2007, 366, 965–975.

[9] D. R. Fuller, C. R. Conant, T. J. El-Baba, C. J. Brown, D. W. Woodall, D. H. Russell, D. E. Clemmer, Journal of the American Chemical Society 2018, 140, 9357–9360.

[10] F. Wang, C. Chen, X. Liu, K. Yang, X. Xu, H. Yang, T. S. Dermody, Journal of Virology 2016, 90, 1910–1917.

[11] L. Zhao, C. D. Kroenke, J. Song, D. Piwnica-Worms, J. J. H. Ackerman, J. J. Neil, NMR in Biomedicine 2008, 21, 159–164.

[12] H. Wang, P. Yang, K. Liu, F. Guo, Y. Zhang, G. Zhang, C. Jiang, Cell Research 2008, 18, 290–301;

18. N. Yang, H.-M. Shen, International Journal of Biological Sciences 2020, 16, 1724–1731.

