## Supporting Information for "Allosteric inhibition of the SARS-CoV-2 main protease – insights from mass spectrometry-based assays"

**Materials and Methods**

*Protein production and purification*

The plasmid encoding the codon-optimized gene for SARS-CoV-2 M^Pro^ fused with an N-terminal glutathione S-transferase tag and a C-terminal His_6_-tag preceded by a human rhinovirus 3C protease site was transformed into E. coli Rosetta (DE3). Several transformed clones were used to inoculate a starter culture which was grown to exponential phase in lysogeny broth at 37 °C containing 100 μg mL^-1^ carbenicillin. The starter culture was used to inoculate 1 L of Terrific Broth Autoinduction media (Formedium) supplemented with 10% (*v*/*v*) glucose and 100 μg mL^-1^ carbenicillin. Cells were grown for 5 h at 37 °C, then cooled to 18 °C and grown for an additional 12 h for recombinant protein production. Cells were harvested by centrifugation (5,000 x g, 15 min).

Cells were resuspended in lysis buffer consisting of 50 mM Tris-HCl (pH 8), 300 mM NaCl, 10 mM imidazole and 0.05 mg/mL benzonase (Sigma Aldrich) and lysed by sonication. Lysates were clarified by centrifugation (50,000 x g, 1 hr). The supernatant was loaded onto a HiTrap Talon Co^2+^ affinity column (GE Healthcare) equilibrated with lysis buffer. The column was washed with lysis buffer supplemented with 25 mM imidazole. Elution was performed with 50 mM Tris pH 8.0, 300 mM NaCl and 500 mM imidazole followed by His_6_-tag cleavage using human rhinovirus (HRV) 3C protease (produced in-house) at 4°C. Reverse immobilized metal affinity chromatography was used to remove the HRV 3C protease and His_6_-tag. The flow-through was concentrated and loaded onto Superdex 75 26/600 size exclusion chromatography column equilibrated with 50 mM Tris-HCl pH 8 and 300 mM NaCl. Protein purity was assessed at each step by SDS-PAGE. Peak fractions for dimeric M^Pro^ were pooled and concentrated before mass spectrometry (MS) analysis.

*11-mer substrate synthesis*

Peptides were synthesized on 0.1 mmol scale as C-terminal amides from C- to N-terminus on a rink amide-MBHA resin (100–200 mesh, 0.6–0.8 mmol g^−1^ loading, AGTC Bioproducts) using a LibertyBlue microwave peptide synthesiser (CEM). *N-α*-Fmoc protected amino acids (CS Bio, Novabiochem, Sigma-Aldrich, TCI, Alfa Aesar, Merck or AGTC Bioproducts); where required acid-labile protecting groups were employed. Coupling and deprotection steps used the instrument’s standard methods and were microwave assisted.^[1]^

*N,N’*-Diisopropylcarbodiimide (TCI Europe) was used as coupling reagent with Oxyma Pure (Merck). For Fmoc deprotection, 20% (v/v) piperidine in DMF (peptide synthesis grade, AGTC Bioproducts) was used. Following completion of coupling reactions, the resin was washed three times with dichloromethane and dried in air. Acid labile protecting groups were removed and cleavage from the resin was conducted using 5 ml of a deprotection mixture: 1,3-cimethoxybenzene (2.5%, 125 ul), triisopropylsilane (TIPS) (2.5%, 125 ul), MilliQ purified water (2.5%, 125 ul) in trifluoroacetic acid (92.5%, 4.625 mL) for 4 h at room temperature. Upon filtration of the resulting mixture, the peptide was precipitated with ice-cold Et_2_O (45 ml). The solid was pelleted (4255 g, 10 min, 4.0 °C); the liquid was decanted off and the solid was dried in air. Before purification, peptides were dissolved in H_2_O and lyophilised to remove trifluoracetic acid. HPLC purification was conducted using a Phenomenex Gemini 250 × 21.2 mm, 110 Å, NX-C18 LC column.

*Sample preparation*

Prior to mass spectrometry (MS) analysis, fresh aliquots of M^Pro^ were buffer exchanged into 200 mM ammonium acetate (pH 7.4) using Zeba spin 7k buffer exchange columns (Thermo Scientific). Fragments were taken from the DSI-poised library as 10 mM stocks in 100% DMSO, diluted with 200 mM ammonium acetate (pH 7.4), and mixed with M^Pro^ in appropriate amounts prior to analysis. For kinetic measurements, 50 µM of 11-mer substrate was mixed with 5 µM of M^Pro^ and analyzed immediately. The inclusion the fragments as well as the 11-mer substrate incorporates 10% DMSO into M^Pro^ containing solutions for MS analysis. Therefore, we have included 10% DMSO in all drug binding and kinetics experiments. All measurements were performed in triplicate.

*Native mass spectrometry*

1–3 µL of protein solution was loaded into in-house prepared gold-coated electrospray capillaries pulled to ~1-3 µm tip diameter.^[^^[2]]^ Samples were analyzed using a Thermo Q-Exactive UHMR Orbitrap platform operated at 30,000 resolving power (at *m*/*z* 200). An electrospray was generated by applying 0.9 – 1.3 kV bias to the electrospray capillary; no backing pressure was applied to the electrospray solution. The instrument capillary was maintained at a temperature of ~100 °C to assist with desolvation. No collision energy was applied throughout the instrument.

*Data Analysis*

To extract the relative populations of each species, mass spectral data were deconvoluted to zero-charge spectra using Unidec software.^[^^[3]]^ Mole fraction for monomer and dimer ratio were calculated at each concentration and fitted to a monomer-dimer binding model described by Bergdoll et al.^[^^[4]]^ Kinetic models were fitted to the experimental data using $\left[ ES \right]_{t}={[ES]}_{0}*e^{-kt}$ where [ES]_t_ and [ES]_0_ are the quantities of enzyme-substrate determined at time *t = t* and *t = 0*, respectively, and *k* is the measured rate constant (in s^-1^). All data models were fit to experimental data sets using a user-defined function in OriginPro 2018 (OriginLab Corporation, Northampton, MA) using a least-squares residual analysis.

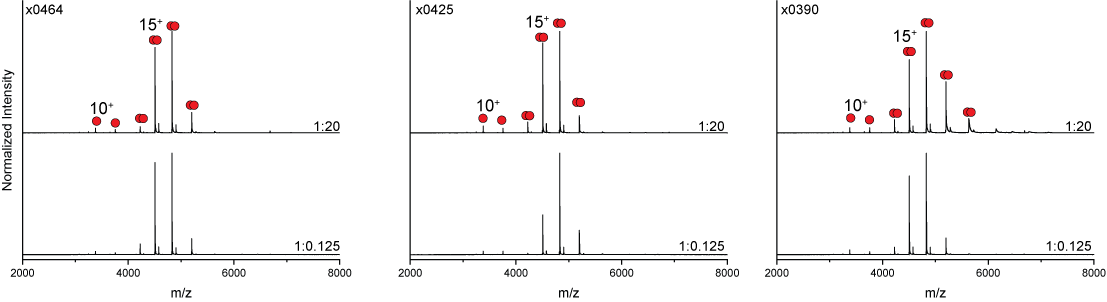

Figure S1. Effect of x0464, x0425, and x0390 on M^Pro^ monomer-dimer equilibria. No substantial differences were observed in the peaks assigned to monomer (10^+^) and dimer (15^+^).

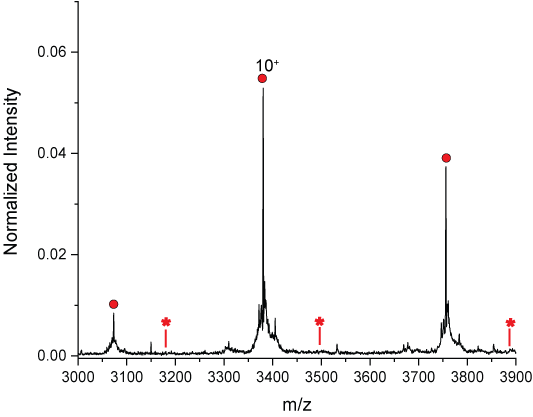

Figure S2. Expansion of the charge state series assigned to monomeric M^pro^ in the presence of 50 mM of the 11-mer substrate 30s after addition of the substrate. The expected peak series for M^Pro^ monomers bound to the 11-mer substrate (indicated with red asterisk) is not observed.

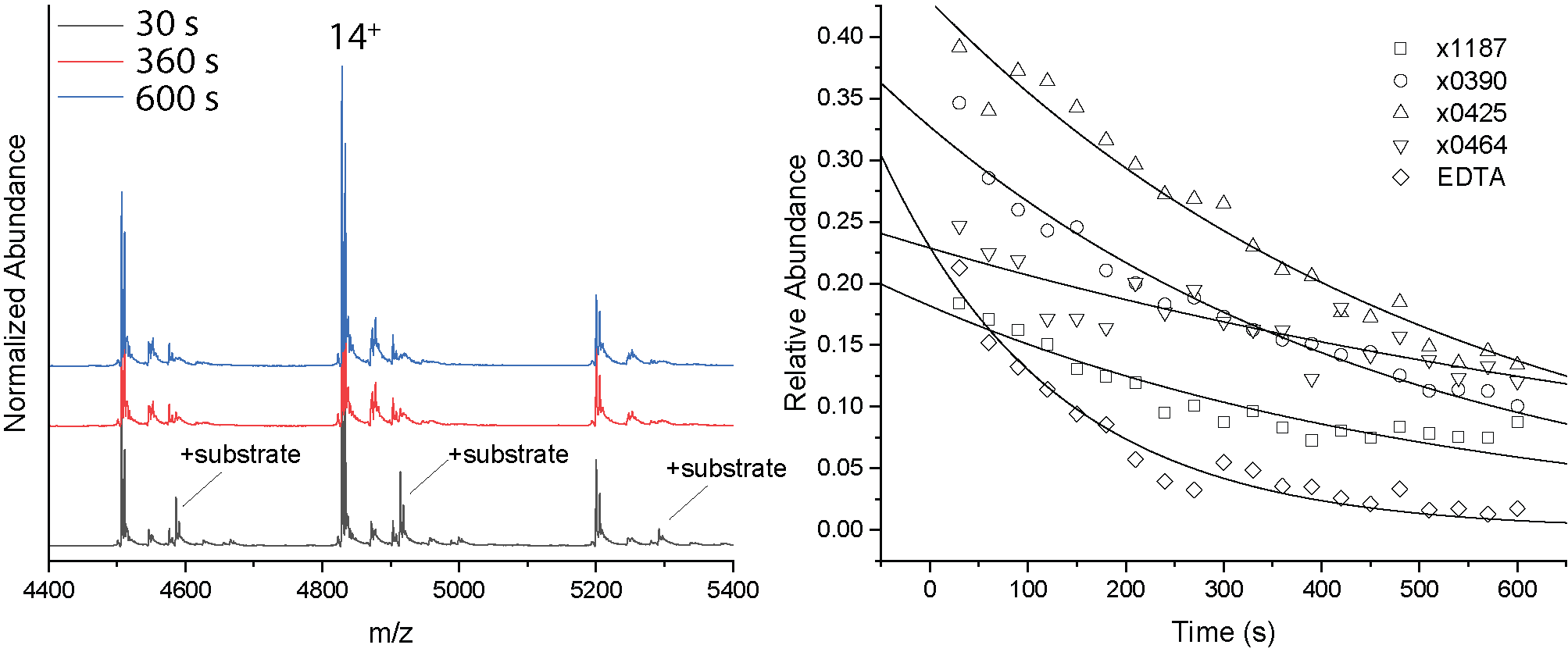

Figure S3. (left) representative kinetics plots showing the degradation of the 50 µM 11-mer substrate by 5 µM M^Pro^ in the presence of 100 µM x1187 (20-fold excess). (Right) representative kinetics plots showing the decay of the enzyme-substrate complex in the presence of 50 µM of the different small molecules. Solid lines are fits to a unimolecular kinetics model to the experimental data (described above).

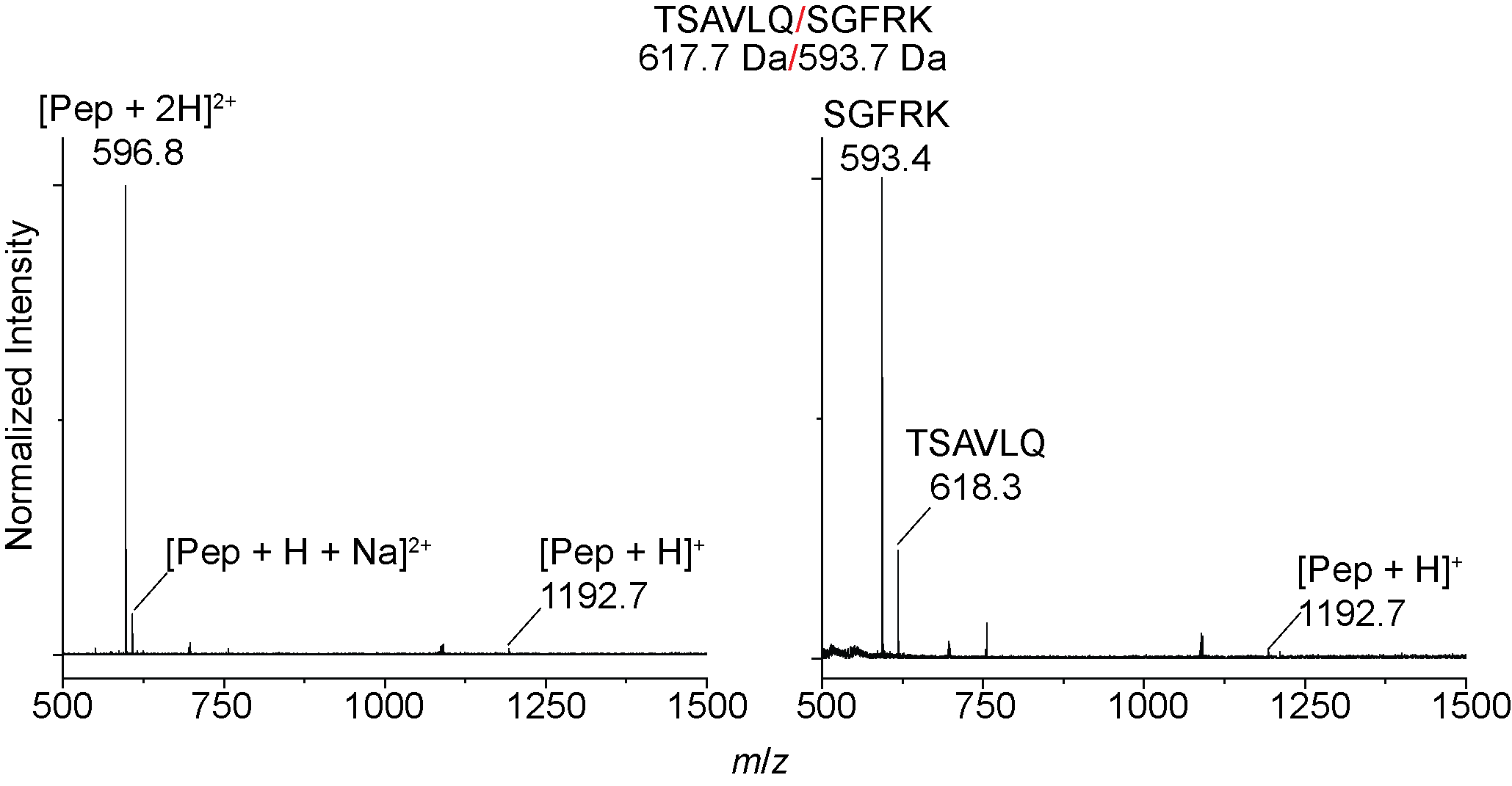

Figure S4. MS data collected for (left) the 11-mer substrate in 200 mM ammonium acetate (pH 7.4) at ~10 µM and (right) the products formed upon M^Pro^ cleavage of the 11-mer substrate at t = 570 s to form two peptide products.

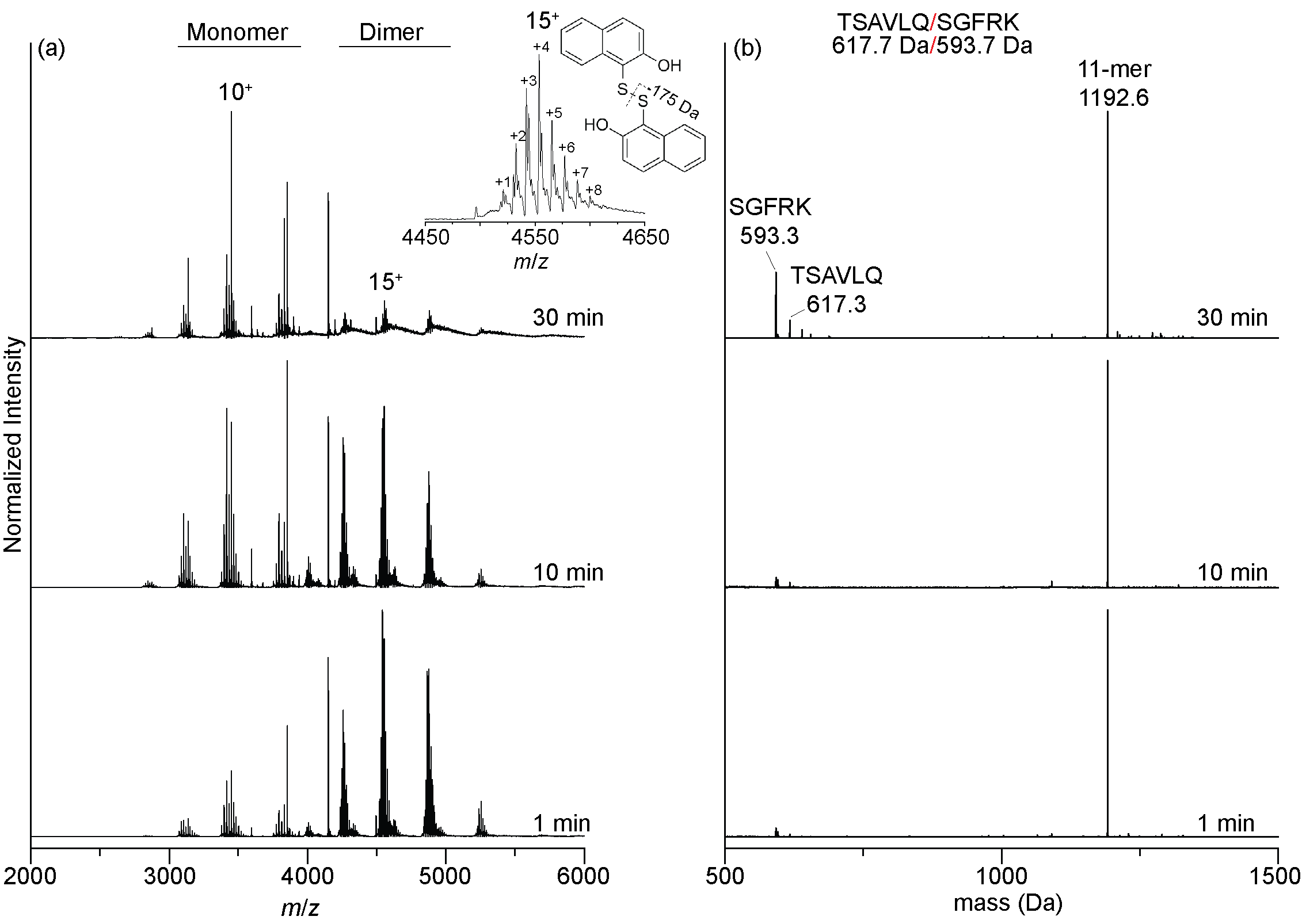

Figure S5. Covalent modification of Cys145 greatly diminishes M^Pro^ proteolytic activity. 5 µM M^Pro^ was mixed with 100 µM IPA3 (20-fold excess prepared in 100% DMSO) and incubated for 15 min. Mass spectra were collected immediately following the addition of 50 µM 11-mer substrate. (a) Native mass spectra collected at several different timepoints showing highly modified M^Pro^ monomers and dimers; no signals for enzyme-substrate complexes were observed. (b) Deconvoluted mass spectra collected under instrument conditions that preferentially transmit low mass-to-charge (m/z) species. At t = 30 min, the dominant feature is the 11-mer substrate – low intensity peaks assigned to the products (m/z 593 Th and m/z 617 Th) are observed.

**References**

1. [] G. W. Langley, M. I. Abboud, C. T. Lohans, C. J. Schofield. *Bioorg. Med. Chem.* **2019**, *27*, 2405-2412.
2. [] H. Hernandez, C. V. Robison. *Nat. Protoc.* **2007**, *2*, 715-726.
3. [] M. T. Marty, A. J. Baldwin, E. G. Marklund, G. K. A. Hochberg, J. L. P. Benesch, C. V. Robinson. *Anal. Chem.* **2015**, *87*, 4370-4376.
4. [] L. A. Bergdoll, M. T. Lerch, J. W. Patrick, K. Belardo, C. Altenbach, P. Bisignano, A. Laganowsky, M. Grabe, W. L. Hubbell, J. Abramson. *Proc. Natl. Acad. Sci. U. S. A.* **2017**, E172-E179.
